# An ancient Hippo–dystrophin axis controls cortical stability and multicellular interactions in a close unicellular relative of animals

**DOI:** 10.64898/2026.11.23.768642

**Authors:** Jonathan E Phillips, Duojia Pan

**Affiliations:** Department of Physiology, Howard Hughes Medical Institute, University of Texas Southwestern Medical Center, Dallas, Texas 75390, USA

## Abstract

How the protein complexes and signal transduction pathways that regulate animal physiology and human disease emerged over evolutionary time remains poorly understood. The dystrophin-dystroglycan complex (DGC), whose disruption causes muscular dystrophy in humans, is best known for its role in maintaining muscle integrity and was previously thought to have originated in animals. Here we show that *Capsaspora owczarzaki*, a close unicellular relative of animals, encodes core DGC components—including a bona fide dystroglycan ortholog—as critical downstream targets of the Hippo pathway. Loss of *Capsaspora* dystrophin (coDystrophin) destabilizes the cortical actin cytoskeleton, causes excessive membrane blebbing, and produces flattened, irregularly shaped multicellular aggregates, all of which phenocopy loss of coYki. Conversely, coDystrophin overexpression rescues the cytoskeletal defects of coYki mutant cells, identifying coDystrophin as a key effector of Hippo signaling.

Despite extensive rearrangement of dystrophin domain architecture during animal evolution, a human microdystrophin partially rescues coDystrophin mutant cells, revealing deep functional conservation. These findings show that core DGC components predate animals and indicate that the complex’s ancestral role was not muscle-specific but lay in maintaining cortical cytoskeletal integrity under the control of an ancient Hippo-regulated genetic module.

## INTRODUCTION

How the functional protein complexes that underlie human health and disease emerged and changed over evolutionary time remains poorly understood. One such complex, the dystrophin-dystroglycan complex (DGC), plays a key role in muscle integrity by coupling the extracellular matrix to the intracellular cytoskeleton and myofibrils via a transmembrane linkage (*1*). At its core, this complex consists of the transmembrane protein dystroglycan, which binds to the extracellular matrix component laminin and interacts directly with the cytoplasmic protein dystrophin (*2, 3*). Although dystrophin has several isoforms that vary in expression by tissue, the largest and most studied isoform (Dp427) is expressed in skeletal and cardiac muscle as well as some classes of neurons (*4, 5*). Dp427 encodes two N-terminal actin-binding calponin homology domains, a long rod-like region of spectrin repeats, and a C-terminal sequence that binds dystroglycan as well as other DGC scaffolding components such as the syntrophins (*6*). The DGC thus serves to couple the ECM to the internal cytoskeleton to maintain the integrity of muscle, further buffering mechanical stress through the “shock-absorbing” properties of the dystrophin rod-like region (*7*). Consistent with this role in muscle function, mutations in dystrophin and other DGC components underlie the muscle wasting disease muscular dystrophy, in which progressive weakness, loss of muscle function, and cardiopulmonary complications occur (*8*). Remarkably, proteins with dystrophin-like domains have been identified in the closest unicellular relatives of animals (*9*). However, genomic analyses of ctenophores, often inferred to represent the earliest-branching animal clade based on current phylogenomic data (*10*), have reported the absence of a dystrophin ortholog (*9, 11*). This fact, and the apparent absence of dystroglycan in unicellular organisms and ctenophores, has been used to argue that the DGC was an animal innovation that emerged after the divergence of ctenophores (*9*).

In order to better understand the evolutionary origins and historical functions of conserved animal signaling pathways and protein complexes, we have developed the unicellular amoeboid organism *Capsaspora owczarzaki* as a genetically tractable research organism (*12*). As a very close unicellular relative of animals, *Capsaspora* possesses many genes and pathways associated with animal physiology, such as an integrin adhesome complex (*13*), tyrosine kinases such as Src and Abl (*14*), a p53/p63/p73 ortholog (*15*), and a cadherin ortholog (*16*). Further, *Capsaspora* cells can aggregate to form spherical multicellular structures stabilized by calcium-dependent cell-cell adhesion (*17*), providing a model for simple multicellularity in a close animal relative. Stable transgenesis and gene knockout techniques have now been established in *Capsaspora*, allowing for functional studies of *Capsaspora* genes (*12*).

We have previously utilized *Capsaspora* to study the premetazoan functions of the Hippo signaling pathway, which was initially discovered for its critical role in restricting tissue growth in animals but later found to be present in both animals and their closest unicellular relatives, such as *Capsaspora* and choanoflagellates (*18, 19*). The Hippo pathway restricts tissue growth by inactivating the transcriptional coactivator Yorkie/YAP/TAZ, a potent driver of cell proliferation in animals (*20, 21*). This regulation occurs through inhibitory phosphorylation of Yorkie/YAP/TAZ mediated by a kinase cascade consisting of the kinases Hippo and Warts ((*22*), Fig. 1a). Besides its role as a potent growth driver in animals, Yorkie/YAP/TAZ also functions as a key intracellular mechanotransducer via which mechanical stimuli, such as cytoskeletal tension and extracellular matrix stiffness, influence gene expression and cellular behaviors (*23*). We found that the *Capsaspora* Yorkie/YAP/TAZ ortholog (coYki) is dispensable for cell proliferation but regulates *Capsaspora* aggregation. Unlike normal *Capsaspora* cells that coalesce into spherical multicellular aggregates, *coYki -/-* cells form flattened, irregular shaped aggregates (*12*). We further traced the defect in aggregate shape to abnormal cortical cytoskeletal dynamics in *coYki -/-* cells: loss of coYki leads to the formation of actin-depleted membrane blebs, and inhibiting membrane blebbing in *coYki -/-* cells rescued the abnormal shape of *coYki* mutant aggregates (*12*). Mutation of the *Capsaspora* Hippo and Warts orthologs (coHpo and coWts) also causes cytoskeletal phenotypes (*24*), further supporting the importance of the Hippo pathway in cytoskeletal regulation in *Capsaspora*. Together, these findings implicate cytoskeletal regulation but not proliferation as an ancestral function of the Hippo pathway.

**Fig. 1:**
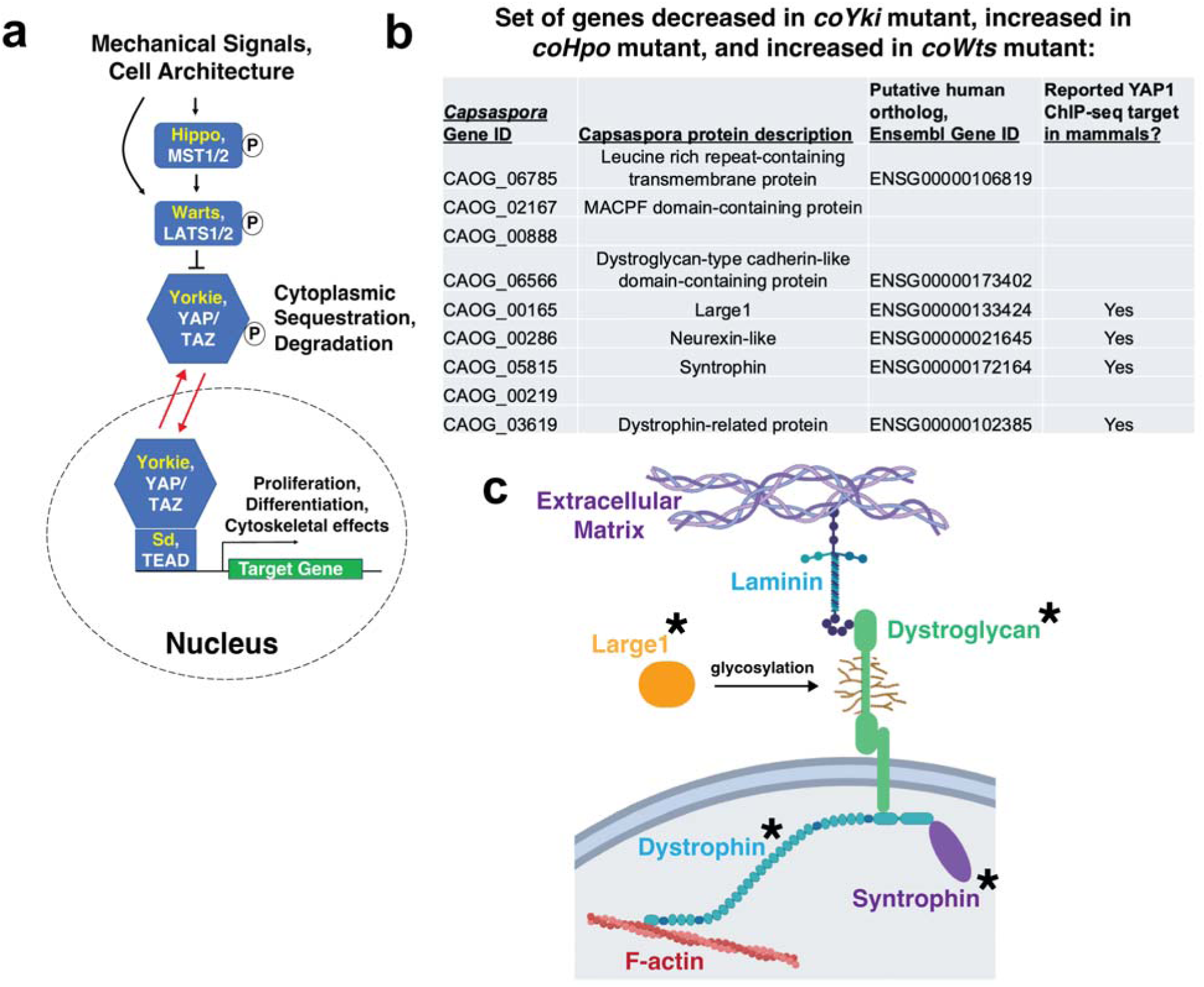
The Hippo pathway regulates the expression of dystrophin-dystroglycan complex genes in the unicellular organism *Capsaspora owczarzaki*. a, Schematic diagram of the Hippo pathway. Names of *Drosophila* orthologs are in Yellow, and mammalian ortholog names are in white. Figure adapted from (*19*). **b,** Using previously obtained RNAseq data (*12, 24*), genes that fit all of the following criteria were selected for: decreased expression in the *coYki* mutant, increased expression in the *coWts* mutant, and increased expression in the *coHpo* mutant. Only nine genes in the *Capsaspora* genome fit this criteria. Putative human orthologs of these *Capsaspora* genes are shown. Absence of a human ortholog means that no ortholog could be assigned (*12*). *Capsaspora* protein descriptions were made based on the assigned protein family and domain annotations from the Uniprot protein database (*25*) for each *Capsaspora* gene ID. Absence of protein description indicates that no relevant data are available. To determine if a human protein is a reported YAP1 ChIP-seq target, 2,212 reported YAP1 ChIP-seq targets were obtained from the Harmonizome 3.0 database (*26*). **c,** Components of the mammalian dystrophin-dystroglycan complex. Asterisk indicates the presence of an orthologous *Capsaspora* gene in the gene list in Fig. 1b.

Despite these advances, how the Hippo pathway mechanistically regulates cytoskeletal dynamics in *Capsaspora* was unknown. In the current study, we attempted to address this open question by identifying putative transcriptional targets of coYki using RNAseq data from *coYki*, *coWts*, and *coHpo* mutant cells. Strikingly, this analysis revealed a selective enrichment in genes linked to the DGC, including putative *Capsaspora* orthologs of dystrophin (coDystrophin) and dystroglycan. We generated a deletion mutant of coDystrophin and found that this mutant phenocopies the cytoskeletal defects of the *coYki* mutant cells. Furthermore, the abnormal cytoskeletal dynamics in *coYki -/-* cells were rescued by overexpression of coDystrophin, highlighting coDystrophin as a key transcriptional target of coYki in cytoskeletal regulation. Our analysis of dystroglycan and dystrophin evolution in unicellular organisms and early-branching animals further revealed that: a) the *Capsaspora* dystrophin ortholog domain structure is representative of the ancestral dystrophin-like protein present in the ancestor of animals, and b) the *Capsaspora* genome encodes a bona fide dystroglycan ortholog, demonstrating that this protein emerged before the origin of animals. Together, these findings identify a direct and deeply evolutionarily conserved role for the Hippo pathway in transcriptional regulation of the DGC, demonstrate that core components of the DGC preceded the origin of animals, and show that regulation of cytoskeletal integrity by dystrophin is a conserved function in a close unicellular relative of animals.

## RESULTS

### The Hippo pathway transcriptionally regulates components of the dystrophin-dystroglycan complex in *Capsaspora*

We previously performed RNAseq on *coYki-/-*, *coHpo -/-*, and *coWts -/-* mutant cells and identified 1205, 1032, and 609 differentially expressed genes compared to WT for each mutant, respectively (*12, 24*). To specifically select for putative targets of coYki, we filtered these datasets for genes that met three criteria: downregulation in the *coYki-/-* mutant, upregulation in the *coHpo -/-* mutant, and upregulation in the *coWts -/-* mutant. Only nine *Capsaspora* genes met these criteria (Fig. 1b). To characterize these genes, we identified their putative mammalian orthologs based on our previous Human-*Capsaspora* orthology analysis (*12*) and the Uniprot Knowledgebase (*25*). Remarkably, four out of the nine putative coYki target genes corresponded to components of the dystrophin-dystroglycan complex (DGC), a membrane-localized protein complex that links the cytoskeleton to the extracellular matrix (Fig. 1c) and whose mutations underlie the muscle-wasting disease muscular dystrophy (*11*). These genes included putative *Capsaspora* orthologs of dystroglycan, a transmembrane receptor, dystrophin, which links dystroglycan to the actin cytoskeleton, syntrophin, which binds dystrophin and recruits signaling molecules such as kinases and ion channels to the complex, and Large1, a Golgi-localized enzyme that glycosylates and is required for the proper function of dystroglycan (*6, 9*)( Fig. 1b,c).

Six of the nine putative coYki target genes have predicted human orthologs. To test if these orthologs may also be YAP1 targets in humans, we assessed whether these genes are present in a set of 2,212 reported YAP1 ChIP-seq targets (*26*). The human orthologs of four of these genes were present in this YAP1 ChIP-seq target set, including dystrophin, syntrophin, and Large1 (Fig. 1b), as well as the dystrophin paralog utrophin. The enrichment of coYki target gene orthologs in this YAP1 ChIP-seq target set is significant (p <0.01, Fisher’s exact test assuming 20,000 human genes). These data indicate that coYki may regulate a gene module associated with the DGC, and that this regulatory module may be ancient and conserved from *Capsaspora* to humans.

### A *Capsaspora* dystrophin mutant phenocopies the *Capsaspora* Yorkie mutant by affecting the shape of multicellular structures

We previously identified two *coYki -/-* mutant phenotypes: aberrant multicellular aggregate morphology and changes in the cortical cytoskeleton that lead to bleb-like protrusions (*12*). If coYki regulates aggregate morphology or the cortical cytoskeleton through transcriptional upregulation of a specific target gene, it is expected that mutation of the target gene may phenocopy the *coYki -/-* mutant. Thus, to test whether coYki may function through any putative coYki target to regulate these cellular processes, we attempted to generate homozygous deletion mutants of these genes using our previously developed gene knockout technique (*12*). We were able to obtain homozygous mutants for CAOG_06785 (*coLRRTM*, for Leucine Rich Repeat TransMembrane), CAOG_00286 (*coNrxn*), CAOG_05815 (*coSyntrophin*), and CAOG_03619 (*coDystrophin*) (Extended Data Fig. 1). We obtained a heterozygous but not homozygous mutant for CAOG_06566 (*coDystroglycan*), suggesting that homozygous mutation of this gene may affect cell viability. To test whether these mutants show aberrant aggregate morphology similar to the *coYki -/-* mutant, we induced cell aggregation and imaged aggregates. *Capsaspora* cells are covered with filopodial projections (Fig. 2a). During aggregation, which can be induced by low-adherence culture conditions, cells adhere to and remain associated with each other through filopodial interactions (Fig. 2b), forming round, three-dimensional structures. *coLRRTM -/-*, *coNrxn -/-*, and *coSyntrophin -/-* aggregates showed a round morphology like WT aggregates (Fig. 2c, e-g). However, like *coYki -/-* aggregates (Fig. 2d), *coDystrophin -/-* aggregates showed an asymmetric, amorphous morphology (Fig. 2h). The similar morphology of *coYki -/-* and *coDystrophin -/-* aggregates suggest that coYki may function through transcriptional regulation of *coDystrophin* to affect cell physiology. Thus we focused on *coDystrophin* for further studies.

**Fig. 2:**
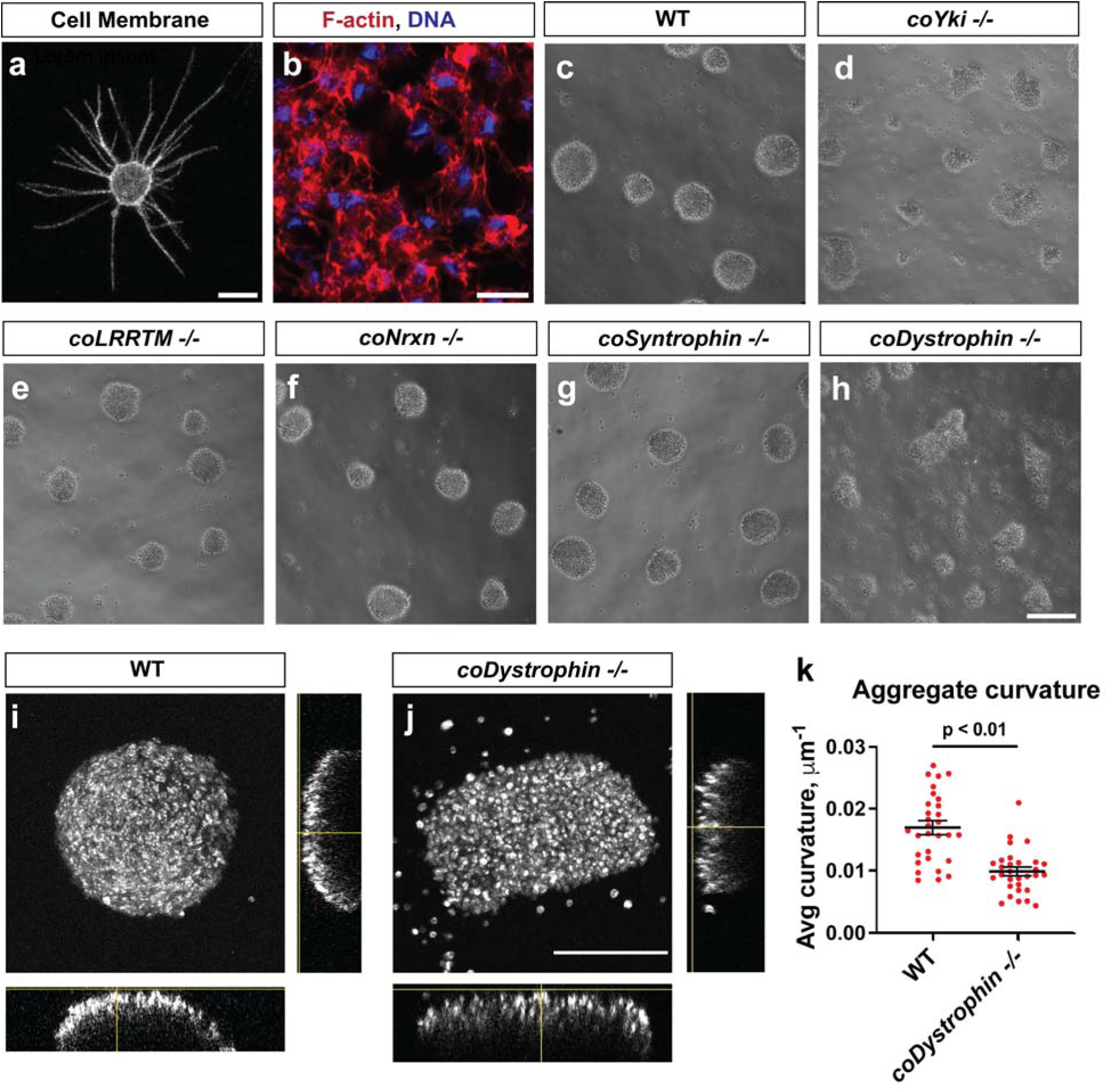
***coDystrophin* mutant cell aggregates resemble *coYki* mutant aggregates. a,** A live *Capsaspora* cell expressing NMM-Venus, a marker of the cell membrane (*53*), which allows visualization of filopodia. Scale bar is 5 microns. **b,** Aggregating *Capsaspora* cells stained with the F-actin marker phalloidin (red) and DNA stain DAPI (blue) to highlight the filopodia-mediated adhesive connections between cells within aggregates. Scale bar is 5 microns. **c-h,** Aggregate morphologies for WT, *coYki* -/-, *coLRRTM* -/- (for <u>L</u>eucine <u>R</u>ich <u>R</u>epeat <u>T</u>rans<u>m</u>embrane, CAOG_06785), *coNrxn* -/- (CAOG_00286), *coSyntrophin* -/- (CAOG_05815), and *coDystrophin* -/- (CAOG_03619) genotypes are shown. Aggregate formation was induced by inoculating cells into low-adherence culture conditions. Scale bar is 75 microns. **i,j,** Live cell aggregates stably expressing a Lifeact-mScarlet transgene were optically sectioned at a region of the aggregate close to the culture chamber surface, and 3D reconstructions were made. A maximum intensity projection of the aggregate and aggregate side views are shown. Scale bar is 75 microns. **k,** Aggregates were imaged as in panels I-J, and the average curvature along the aggregate surface from side view reconstructions was calculated using Kappa (*48*). Average curvature is in units 1/R, where R is the radius of curvature at a given point. Red dots represent measurements from individual aggregates, and black bars indicate mean ± SEM of the mean curvature from 3 independent experiments. The difference between WT and *coDystrophin -/-* is significant (p < 0.01, t-test).

coYki affects the three-dimensional shape of cell aggregates: whereas WT aggregates exhibit a spheroid morphology, *coYki -/-* aggregates are flatter and remain closely apposed to the surface on which they develop (*12*). To test whether *coDystrophin* loss produces a similar phenotype, we optically sectioned live *coDystrophin -/-* aggregates to generate three-dimensional reconstructions of aggregate shape. As previously observed, WT aggregates showed a round, spheroid morphology (Fig. 2i,k). In contrast, *coDystrophin -/-* aggregates were flatter and showed a reduced curvature as compared to WT (Fig. 2j,k), highlighting further similarities between the *coYki* and *coDystrophin* mutants.

To confirm that the aberrant multicellular aggregate morphology of the *coDystrophin -/-* cell line is specifically due to the loss of function of the *coDystrophin* gene, we attempted to rescue this phenotype by expressing a *mScarlet-coDystrophin* fusion transgene. In animals, Dystrophin localizes to the cytoplasmic face of the cell membrane in skeletal muscle (*27*). To examine the localization of mScarlet-coDystrophin, we stably expressed this fusion protein or mScarlet as a control in *Capsaspora* cells. Whereas mScarlet was localized throughout the cytoplasm, mScarlet-coDystrophin was enriched at the cell membrane (Fig. 3a), suggesting that mScarlet-coDystrophin was properly localized in *Capsaspora* cells. We then asked whether mScarlet-coDystrophin expression could rescue the phenotype of *coDystrophin -/-* cells. We found that, when mScarlet-coDystrophin was stably expressed in the *coDystrophin -/-* mutant, aggregate morphology resembled WT (Fig. 3b). Quantification of aggregate size and shape showed that aggregate size was not significantly different between genotypes (Fig. 3c), but that aggregate non-circularity was significantly increased in *coDystrophin -/-* aggregates, and that this increase was rescued by expression of mScarlet-coDystrophin in the mutant line (Fig. 3d). These results indicate that the aberrant aggregate morphology phenotype of the *coDystrophin -/-* mutant is specifically due to loss of function of the *coDystrophin* gene.

**Fig. 3:**
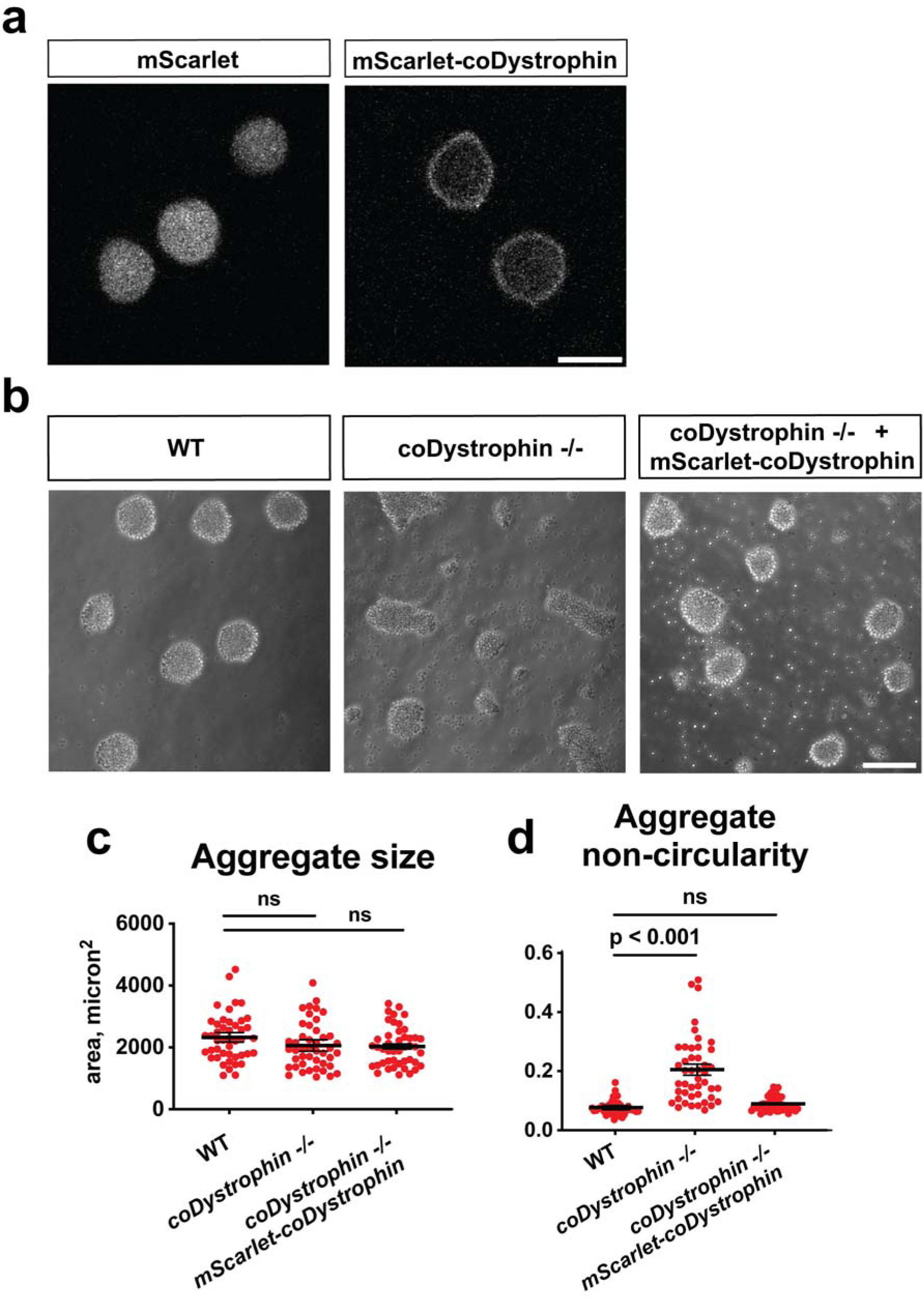
Genetic rescue of the *coDystrophin -/-* aggregation phenotype. a, Live *Capsaspora* cells stably expressing either mScarlet or a mScarlet-coDystrophin fusion protein. mScarlet is localized throughout the cytoplasm, whereas mScarlet-coDystrophin is enriched at the cell periphery. Scale bar is 5 microns. **b,** Aggregation was induced by inoculating cells into low-adherence conditions. Expression of mScarlet-coDystrophin rescues the aggregation phenotype of *coDystrophin -/-* cells. Scale bar is 75 microns. **c,** Aggregate size was quantified by manual tracing using ImageJ. Three independent experiments were done, with 15 aggregates per genotype measured for each replicate. Red dots represent individual aggregates, and black bars show mean ± SEM of the mean aggregate sizes from each independent experiment. Differences between genotypes are not significant (one-way ANOVA, Dunnett’s test). **d,** Aggregate non-circularity was quantified by manual tracing of aggregates. ImageJ was used to quantify circularity, and non-circularity was calculated using the formula (1 – circularity). Three independent experiments were done, with 15 aggregates measured for each experiment. Red dots represent individual aggregates, and black bars show mean ± SEM of the mean non-circularity from each independent experiment. Absence of error bars indicates that error is smaller than the plot symbol. The difference in circularity is significant between WT and *coDystrophin -/-* (p < 0.001), but not between WT and the *coDystrophin -/- mScarlet-coDystrophin* rescue cell line (one-way ANOVA, Dunnett’s test).

### The domain architecture of *Capsaspora* Dystrophin is representative of an ancestral Dystrophin protein

Despite the sequence homology between coDystrophin and human dystrophin, their domain organizations differ in significant ways. Human dystrophin includes two actin-binding calponin homology domains (CHDs) at its N terminus, followed by a long, rod-like region of 24 spectrin-like domains that has been proposed to act as a flexible, shock-absorbing structure during muscle contraction (*7*) (Fig. 4a). C-terminal to these spectrin repeats is a multi-domain module specific to dystrophin-related proteins consisting of a WW domain, an EF hand domain, and a ZZ-type zinc-finger domain -- we will hereafter refer to this multi-domain module as the <u>c</u>ysteine-rich <u>d</u>omain (CRD), a name previously used (*28, 29*). The CRD is necessary for interaction with Dystroglycan, in part through binding of the dystrophin WW domain to a PPxY motif at the intracellular C-terminus of Dystroglycan (*29, 30*). Compared to the human dystrophin, coDystrophin lacks any spectrin repeats. It also contains a CRD and a CHD in a reversed configuration, with a CRD at its N-terminus and a CHD at its C-terminus (Fig. 4a).

**Fig. 4:**
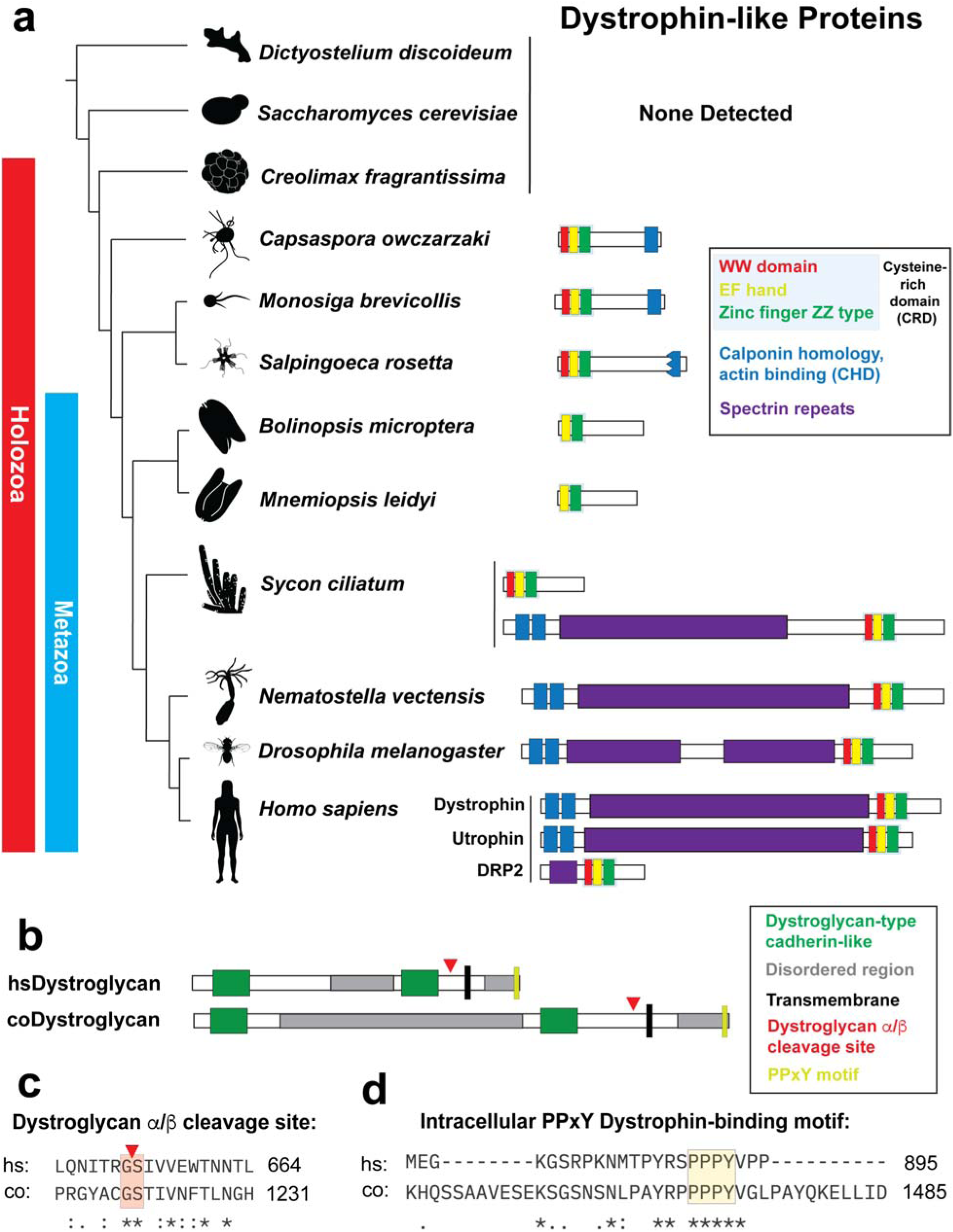
Evolution and conservation of Dystrophin and Dystroglycan. a, Dystrophin-like proteins from different species are shown. “Dystrophin-like protein” indicates a protein that contains the WW domain, EF Hand, and ZZ type domain module (i.e., the Cysteine-rich domain, or CRD) present in mammalian dystrophin. Also included are proteins from *Bolinopsis microptera* and *Mnemiopsis leidyi*, in which the WW domain has apparently been lost. For the displayed species, all proteins in which the CRD was found are shown. The dystrophin-like protein from *Salpingoeca rosetta* contains a truncated CHD (indicated by jagged edge). **b,** The structure of human Dystroglycan (hsDystroglycan) and the *Capsaspora* Dystroglycan ortholog (coDystroglycan). **c,** Alignment of the human and *Capsaspora* Dystroglycan sequences shows conservation of the “Glycine-Serine” site where autoproteolytic cleavage generates the Dystroglycan α and β subunits. Red triangle indicates cleavage site. **d,** The intracellular PPxY motif at the C-terminus of Dystroglycan that binds the WW domain in dystrophin is conserved in the *Capsaspora* Dystroglycan ortholog.

The difference in domain organization between human dystrophin and coDystrophin could be due to two historical possibilities: The coDystrophin domain architecture could represent an ancient, ancestral form of dystrophin-like protein from which the human dystrophin evolved, or coDystrophin could represent a more recently evolved domain architecture not representative of the ancestral protein. To evaluate these possibilities, and to better understand dystrophin evolution in early animals and close unicellular relatives, we examined domain architectures of dystrophin-related proteins in early-branching animals and unicellular organisms (Fig. 4a). No dystrophin-like proteins (as characterized by the presence of the CRD) were detected in amoebozoa, fungi, or ichthyosporea. coDystrophin is the most basal dystrophin-like protein detected, suggesting that this protein family originated in a unicellular ancestor of *Capsaspora* that gave rise to the animal lineage. Dystrophin-like proteins were also detected in the choanoflagellates (the closest unicellular relatives of animals) *S. rosetta* and *M. brevicollis*. These choanoflagellate dystrophin-like proteins showed a domain organization like coDystrophin, although the CHD in *S. rosetta* is truncated. The fact that this domain organization is shared between *Capsaspora* and choanoflagellates implicates it as an ancient (at least as old as the common ancestor of these two lineages) and likely ancestral form of dystrophin-like proteins.

Current genomic evidence suggests that ctenophores are the earliest branching animal group (*10*). We identified dystrophin-like proteins in the ctenophores *Bolinopsis microptera* and *Mnemiopsis leidyi,* although these ctenophore proteins lack the WW domain within the CRD and do not possess a CHD. Like their *Capsaspora* and choanoflagellate counterpart, the CRD in these ctenophore orthologs is at their N-terminus, and no spectrin repeats are present (Fig. 4a). Thus the domain organization of the ctenophore dystrophin-like proteins more closely resembles the unicellular dystrophin-like proteins than human dystrophin. These data reveal that dystrophin-like proteins in the earliest animals were more similar in structure to coDystrophin than to human dystrophin.

In sponges, another early-branching animal clade, we detected a species (*Sycon ciliatum*) with two paralogs of dystrophin-related proteins: one with an N-terminal CRD as in coDystrophin, and another with a domain organization very similar to human dystrophin (N-terminal CHDs, spectrin repeats, and a C-terminal CRD). In species from later-branching lineages, such as the cnidarian *Nematostella* or *Drosophila,* only dystrophin-related proteins with the human dystrophin architecture were detected. These data suggest that, in an animal ancestor of sponges and humans, gene duplication and paralog evolution led to two dystrophin-like proteins: one resembling coDystrophin with N-terminal CRD, and another resembling human dystrophin with C-terminal CRD. Both paralogs are present in sponges, but the paralog similar to coDystrophin was subsequently lost in the lineage leading to humans. Further gene duplications resulted in the human dystrophin paralogs utrophin and DRP2. Remarkably, despite their considerable difference in domain architecture and evolutionary distance, coDystrophin and human dystrophin both encode a CRD and at least one CHD, and therefore could both plausibly perform a similar function: to link the cytoskeleton to the extracellular environment through dystroglycan.

### The *Capsaspora* genome encodes a Dystroglycan ortholog

A previous analysis reported that the transmembrane protein dystroglycan originated in the animal lineage and is not present in unicellular organisms such as choanoflagellates and *Capsaspora* (*9*). Because our orthology analysis revealed that one of our putative coYki targets (CAOG_06566) showed homology to dystroglycan, we examined sequence similarities between this putative coDystroglycan and human dystroglycan. Both proteins encode an extracellular domain consisting of two dystroglycan-type cadherin-like domains separated by a disordered region, a transmembrane sequence, and an intracellular disordered region (Fig. 4b). Human dystrogycan is autoproteolytically cleaved into an extracellular α subunit and a membrane-spanning β subunit, which stay associated though noncovalent interactions after cleavage (*31*). In human dystroglycan this cleavage occurs at the GS motif at G653 (*32*). Alignment of human dystroglycan and coDystroglycan showed conservation of this GS motif, which is located near the N-terminal side of the transmembrane segment in both proteins (Fig. 4b,c). Human dystroglycan binds the WW domain of dystrophin via a PPxY motif found at the C-terminal end of the β subunit (*30*). We observed conservation of this PPxY motif and surrounding residues at the C-terminus of coDystroglycan (Fig. 4d). Similarities in domain organization, conservation of the α/β cleavage site, and conservation of the C-terminal dystrophin binding PPxY motif strongly suggest that coDystroglycan is a true dystroglycan ortholog. These findings highlight the ancient, premetazoan origins of the DGC.

### Phenotypic suppression by blebbistatin indicates a common underlying mechanism for Dystrophin and Yorkie mutant aggregate phenotypes

Although *coYki -/-* and *coDystrophin -/-* mutant aggregates display a similar aberrant morphology, it is formally possible that aggregate morphology could be perturbed in these mutants for different underlying reasons. For example, defects in cell adhesion, filopodial function, or cell motility could potentially affect aggregate formation. We thus sought to test whether similar causes underlie the aggregate morphology phenotype of these two mutants, which would further support the model that coYki functions through transcriptional regulation of coDystrophin to affect cell physiology. We previously showed that the myosin II inhibitor blebbistatin not only suppressed the excessive blebbing in *coYki -/-* mutant cells but also rescued the aberrant morphology of *coYki -/-* aggregates, suggesting a causal link between cytoskeletal defects and aggregate morphology (*12*). To test whether blebbistatin can similarly rescue aberrant aggregation in the *coDystrophin -/-* mutant, we induced cell aggregation in the presence or absence of blebbistatin. As a control for the specificity of response to blebbistatin, we assayed the effect of blebbistatin on aggregation of the *coWts -/-* mutant, which also shows aberrant aggregation but in a manner distinct from the *coYki -/-* mutant (*24*). As previously reported (*12*), blebbistatin had no effect on the morphology of WT aggregates but suppressed the aberrant aggregate morphology of *coYki -/-* aggregates as assayed by aggregate non-circularity (Fig. 5). Blebbistatin did not suppress the aberrant aggregate morphology of the *coWts -/-* mutant, demonstrating the specificity of the effect of blebbistatin on aggregate morphology. Strikingly, blebbistatin also suppressed the aberrant aggregate morphology of the *coDystrophin -/-* mutant. These results suggest that similar underlying causes are responsible for the abnormal aggregate phenotype in the *coYki -/-* and *coDystrophin -/-* mutants, further supporting *coDystrophin* as a key transcription target of coYki in regulating cell physiology.

**Fig. 5:**
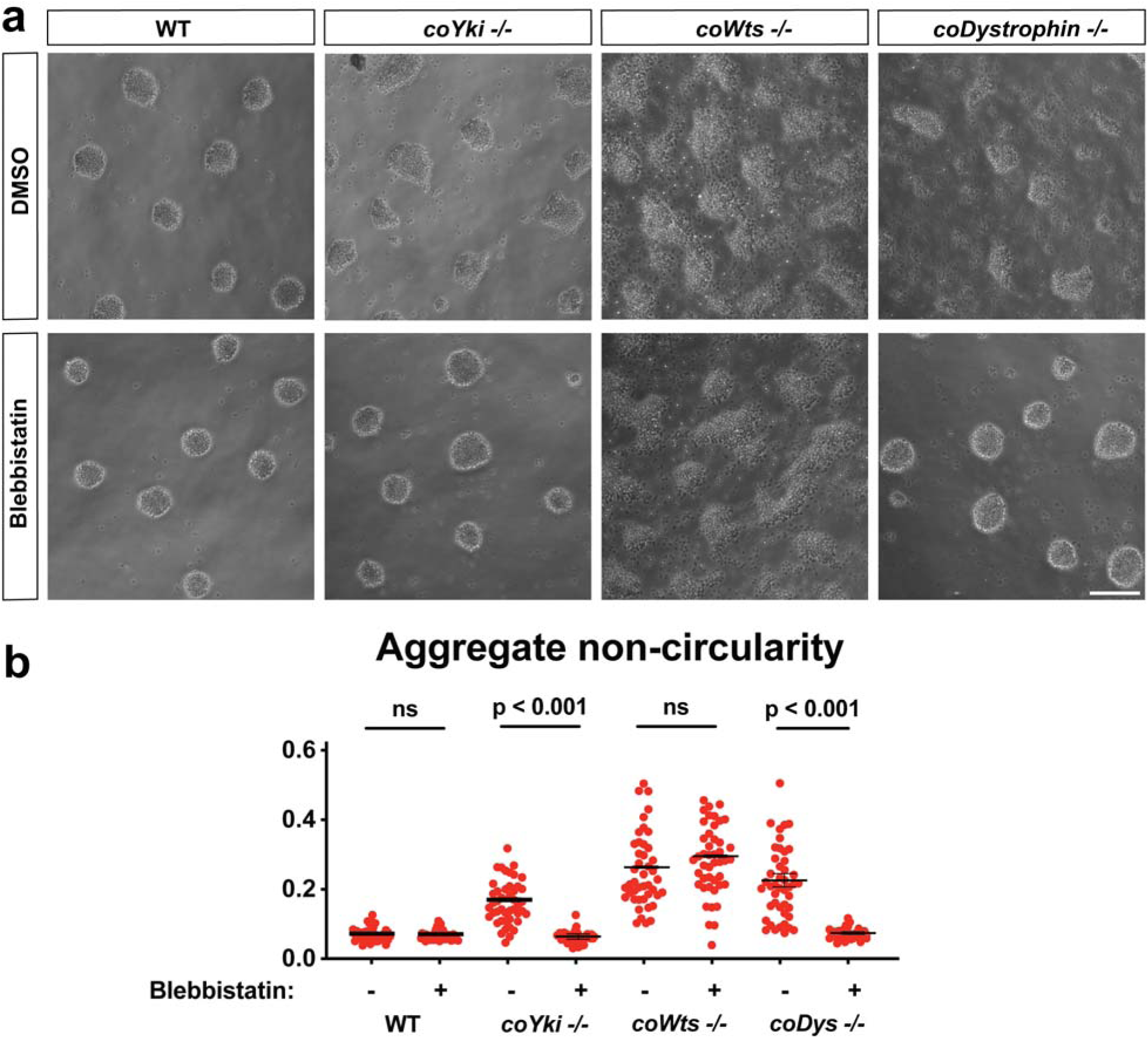
Like the *coYki* mutant, the *coDystrophin* mutant aggregate phenotype can be suppressed by blebbistatin. a, Cells in growth medium containing either DMSO (vehicle control) or 1 μM blebbistatin were inoculated into low-adherence conditions to induce aggregation. Scale bar is 75 microns. **b,** Imaged aggregates were manually traced, the circularity of aggregates was quantified using ImageJ, and non-circularity was calculated using the formula (1-circularity). Red dots indicate measurement values for individual aggregates, and black bars indicate mean ± SEM of the mean non-circularity from each independent experiment (n = 3). Absence of error bars indicates that error is smaller than the plot symbol. For *coYki* and *coDystrophin* mutants, the differences in non-circularity with or without blebbistatin are significant (p < 0.001, one-way ANOVA, Tukey’s test).

### Dystrophin enhances the integrity of the cortical cytoskeleton in *Capsaspora*

In addition to the aberrant aggregate morphology phenotype, *coYki -/-* mutant cells also display a cytoskeletal phenotype. WT *Capsaspora* cells are motile, but show a stable circular F-actin cortex at the periphery of the cell body. In contrast, *coYki -/-* cells show asymmetric distribution of F-actin at the cell cortex. When examined by time lapse microscopy, these regions of F-actin asymmetry in the *coYki -/-* mutant correlate with actin-depleted dynamic protrusions at the cell surface that resemble membrane blebs in animal cells (*12*). These F-actin-depleted protrusions then accumulate F-actin at the distal end and contract back into the cell body, resulting in the appearance of hyperactive cytoskeletal dynamics at the cortex. To test whether *coDystrophin -/-* cells show similar bleb-like protrusions, we used time-lapse microscopy to image live cells expressing the F-actin marker Lifeact-mScarlet. As previously observed, WT cells displayed motility but consistently showed enrichment of F-actin at a uniform round cell cortex (Fig. 6a,b and Video S1). In contrast, *coDystrophin -/-* cells showed asymmetric F-actin distribution at the cell cortex, with dynamic bleb-like actin-depleted protrusions visible by time-lapse microscopy (Fig. 6a,b and Video S2). Quantification of the frequency of these blebs showed that they were very rare in WT cells, whereas they appeared at a rate of approximately two per minute in *coDystrophin -/-* cells, similar to the rate of three per minute observed in *coYki -/-* cells (*12*). These findings reveal similar functions of coYki and coDystrophin in stabilizing the cortical cytoskeleton, further strengthening our model implicating *coDystrophin* as a key transcriptional target of coYki.

**Fig. 6:**
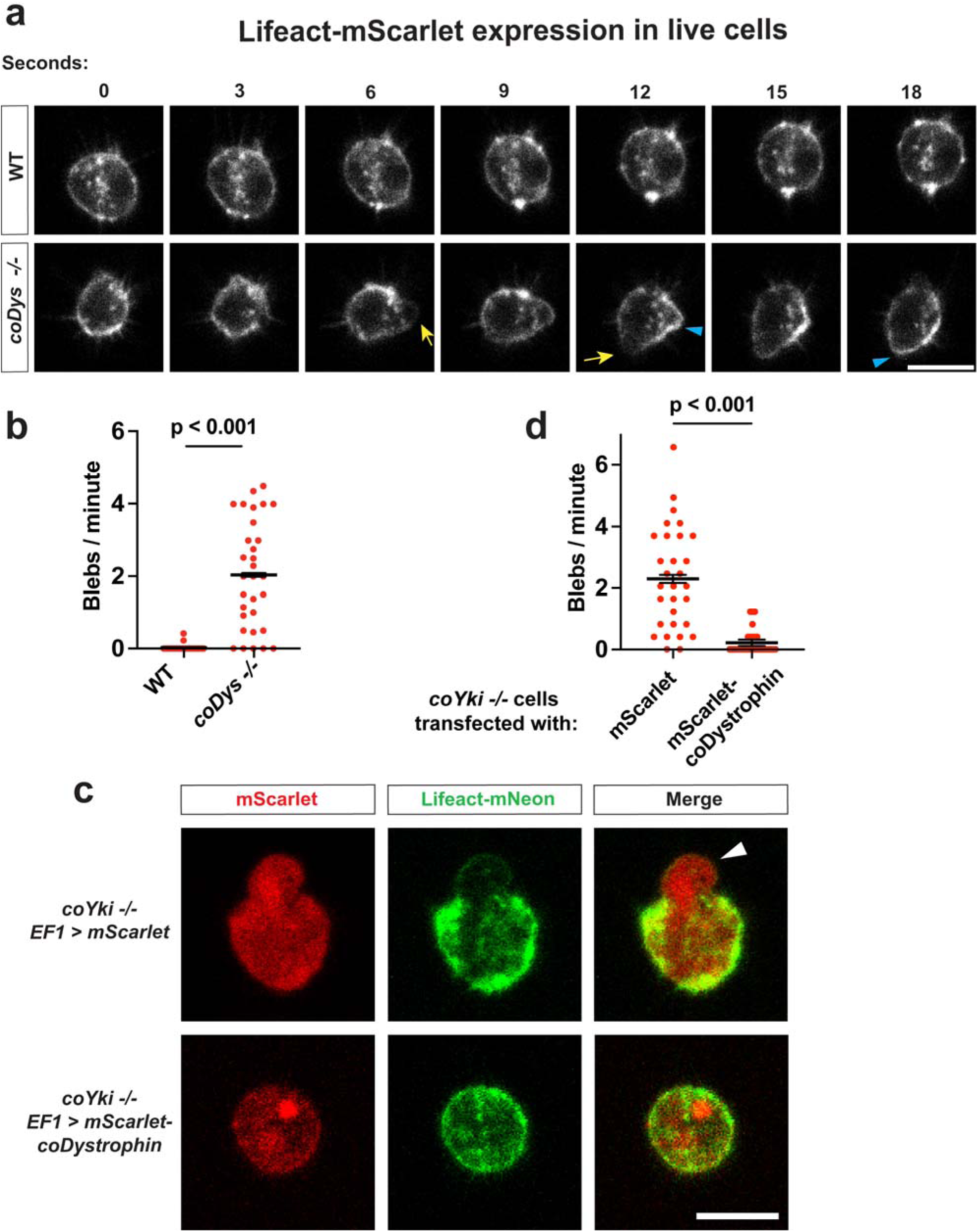
***coDystrophin* stabilizes the cortical cytoskeleton in *Capsaspora*. a,** Live adherent cells stably expressing the F-actin marker Lifeact-mScarlet were imaged by time-lapse confocal microscopy. Yellow arrows indicate F-actin-depleted cell protrusions (“blebs”), and blue triangles indicate F-actin-enriched regions of the cell periphery contracting back to the cell body. Scale bar is 5 microns. **b,** The rate of bleb formation in WT and *coDystrophin -/-* cells was quantified. Cells were imaged as in panel A and the number of blebs per time interval was counted. Red dots indicate calculated blebs/minute values for individual cells, and black bars show mean ± SEM of the mean blebs/minute values from each independent experiment (n=3 with 10 cells measured for each independent experiment). Absence of error bars indicates that error is smaller than the plot symbol. The difference in blebs/minute between WT and *coDystrophin -/-* cells is significant (t-test). **c,** *coYki* -/- cells were transiently transfected with transgenes encoding Lifeact-mNeon and either mScarlet or an mScarlet-coDystrophin fusion protein. Cells were then imaged live by confocal microscopy. Arrowhead indicates an F-actin-depleted bleb. Scale bar is 5 microns. **d,** The rate of bleb formation in *coYki -/-* cells transiently transfected with the indicated transgenes and Lifeact-mNeon was quantified as described for panel B. The difference in blebs/minute between mScarlet and mScarlet-coDystrophin conditions is significant (t-test).

If coYki positively regulates *coDystrophin* expression, and if the aberrant cytoskeletal properties in *coYki -/-* cells are primarily due to reduced *coDystrophin* levels, ectopic *coDystrophin* expression may rescue these cytoskeletal defects. To test this prediction, we transiently transfected *coYki -/-* cells with *coDystrophin-mScarlet* and *Lifeact-mNeon* transgenes simultaneously to visualize the cytoskeletal effects of increased *coDystrophin* expression. When *mScarlet* was transiently expressed as a control, *coYki -/-* cells showed blebbing as previously observed (Fig. 6c,d and Video S3 (*12*)). However, when coDystrophin-mScarlet was transiently expressed, *coYki -/-* cells showed a stable actin cortex, and blebbing was sharply reduced (Fig. 6c,d and Video S3). Thus, the increased stability of the actin cortex in WT cells as compared to *coYki -/-* cells is to a large extent due to positive transcriptional regulation of *coDystrophin* by coYki. This result further supports a role for *coDystrophin* in stabilizing the actin cortex and implicates *coDystrophin* as a key transcriptional target of coYki.

### Conservation of function in Human and *Capsaspora* Dystrophin orthologs

Although human dystrophin and coDystrophin differ considerably in domain organization, both contain a CHD and a CRD, which bind to actin and to dystroglycan, respectively (*6*). If these two activities underlie coDystrophin’s role in cytoskeletal regulation, then human Dystrophin might similarly substitute for this function in *Capsaspora*. To test this possibility, we expressed human dystrophin (hsDystrophin) in *coDystrophin -/-* cells and tested whether it could rescue the blebbing phenotype. To overcome the difficulties of expressing an enormous protein such as human dystrophin (427 kDa), we utilized microdystrophin (138 kDa), a truncated engineered form of human dystrophin developed to facilitate gene therapy applications (*33*). We found that hsDystrophin could indeed rescue the blebbing phenotype of the *coDystrophin* mutant, although the degree of rescue was less than that of coDystrophin expression (Fig. 7 and Video S4). These results support functional conservation between human and *Capsaspora* dystrophin orthologs despite the different relative position of the CHD and the CRD within these proteins.

**Fig. 7:**
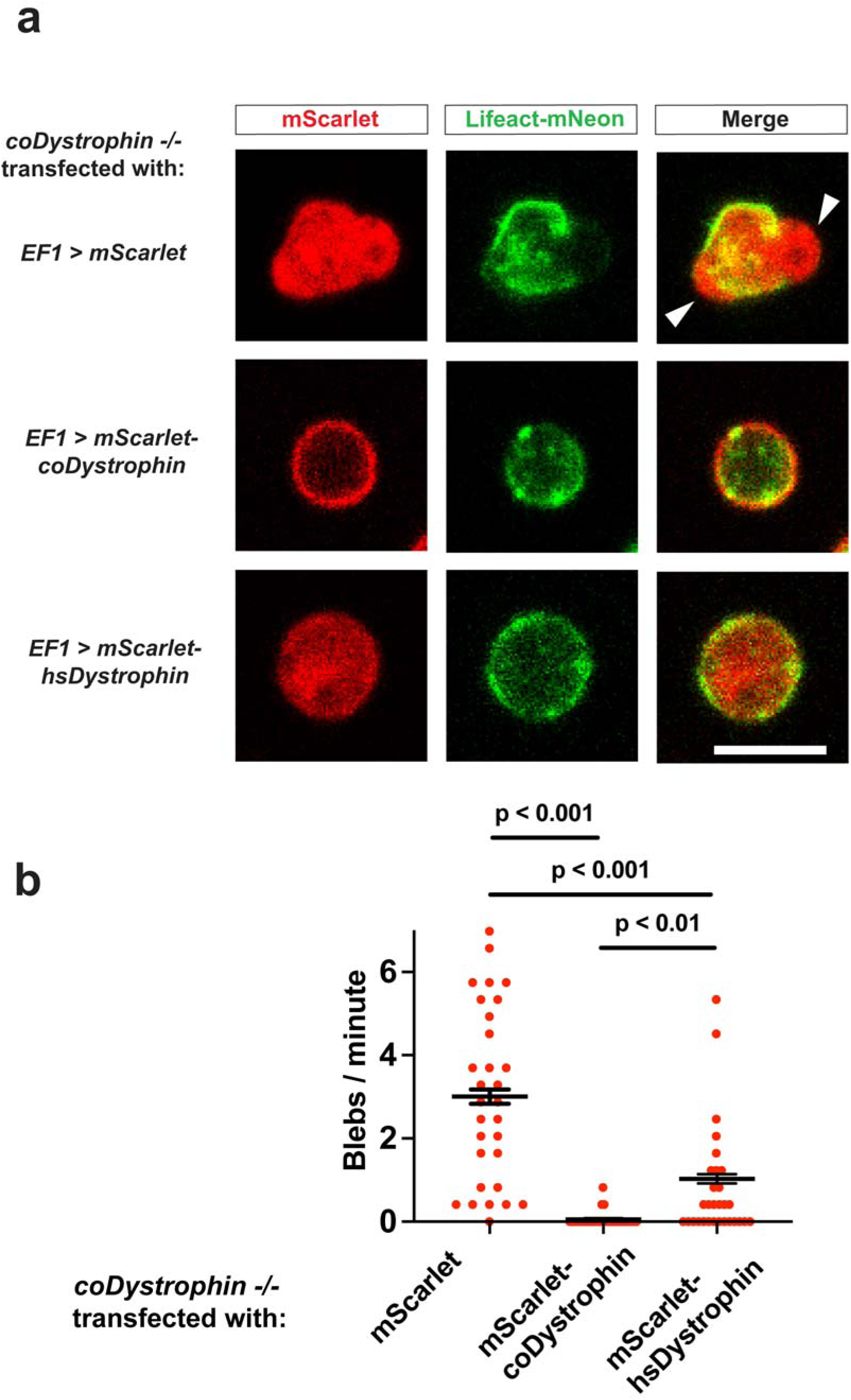
Human Dystrophin rescues the *coDystrophin -/-* blebbing phenotype. a, *coDystrophin* -/- cells were transiently transfected with Lifeact-mNeon and either mScarlet, mScarlet-coDystrophin, or mScarlet-hsDystrophin. Cells were then imaged by confocal microscopy. Arrowheads indicate actin-depleted blebs. Scale bar is 5 microns. **b,** The rate of bleb formation was quantified for *coDystrophin -/-* cells transiently transfected with the indicated constructs using time lapse microscopy. Red dots indicate measurements from individual cells, and black bars indicate the mean ± SEM of the calculated means from each independent experiment. Absence of error bars indicates that error is smaller than the plot symbol. The indicated differences are significant (one-way ANOVA, Tukey’s test).

## Discussion

Much remains to be understood about how the conserved pathways and complexes that underlie animal physiology and human disease emerged over evolutionary time. Rather than viewing these systems only through their modern functions, reconstructing their ancestral roles can reveal the cellular processes they originally evolved to mediate and enhance our understanding of their current functions. Here, we show that the dystrophin-dystroglycan complex (DGC), a key mediator of muscle integrity and cell-ECM interactions in animals, predates the origin of animals and functions in *Capsaspora owczarzaki* as part of a Hippo-regulated program that maintains cortical cytoskeletal integrity. These findings reveal a premetazoan Hippo–DGC regulatory module and suggest that the DGC did not originate as a muscle-specific apparatus, but instead emerged before the origin of muscle as a regulator of cortical integrity.

We found that components of the DGC were strongly enriched among a stringent set of candidate coYki transcriptional targets, defined by reciprocal regulation in *coYki*, *coHpo*, and *coWts* mutant RNAseq datasets. Importantly, our functional studies implicate *Capsaspora* dystrophin ortholog (coDystrophin) as a key transcriptional target of the Hippo pathway. To our knowledge, dystrophin has not previously been highlighted as a YAP/TAZ/Yorkie regulatory target. However, both dystrophin and its mammalian paralog utrophin were identified as YAP-bound genes in a ChIP-seq analysis in mouse embryonic stem cells (*34*). Further, sarcoglycan delta, a component of the mammalian DGC, was identified as a YAP target by ChIP-seq in regenerating mouse heart (*35*) although the DGC was suggested to play an opposing role to YAP in this context (*36*). Further studies will be required to determine whether Hippo-dependent regulation of the DGC contributes to cortical integrity, cytoskeletal organization, or mechanotransduction in embryonic stem cells, regenerating heart, or other animal systems.

Previous reports have suggested that dystrophin and dystroglycan are present in early-branching animals such as sponges but not in unicellular organisms, indicating a metazoan origin of the DGC (*9, 11*). Our identification of a candidate unicellular dystroglycan ortholog and evidence of functional conservation between *Capsaspora* and human dystrophin strongly argue for a premetazoan origin of this complex, highlighting a deeply conserved role for the DGC in cortical stability. The very similar domain structures of dystrophin orthologs from the deeply diverged unicellular lineages of *Capsaspora* and choanoflagellates suggest that this domain architecture was already present in their common ancestor and likely represents the structure of the ancestral dystrophin ortholog. However, extensive rearrangement occurred during animal evolution: whereas unicellular orthologs have a single C-terminal actin-binding CH domain, mammalian dystrophin has two N-terminal CH domains and a long internal region of spectrin repeats that is absent from unicellular and some early-branching animal orthologs. Interestingly, this “shock absorbing” (*7*) spectrin repeat region is found in some sponge species (Fig. 4), revealing that this stress-mitigating domain structure emerged before the emergence of true muscle tissue (*37*). Further, our analysis shows that some early-branching animal dystrophin orthologs have lost the canonical WW domain, and thus are predicted to have reduced or ablated binding to dystroglycan (*38*). Thus dystrophin has undergone extensive structural diversification during animal evolution, making our observed functional conservation of dystrophin from *Capsaspora* to humans particularly striking.

Given the well-established role of the DGC in cell-ECM interactions in animals (*1*), our identification of a critical role for coDystrophin in *Capsaspora* aggregation highlights the possibility of a significant contribution by cell-ECM interactions in the emergence of animal multicellularity. This inference is also supported by studies showing that the ECM-associated protein Rosetteless is required for multicellular rosette formation in the choanoflagellate *S. rosetta*, another close unicellular relative of animals (*39*). Interestingly, although *Capsaspora* encodes a single cadherin-like transmembrane protein, its knockout does not affect the aggregation process or the morphology of aggregates (our unpublished data). We speculate that cell-ECM interactions played a dominant role in the physical connection of cells in the transition to multicellarity and preceded the functional co-option of cadherin-like molecules into cell-cell adhesion proteins. *Capsaspora* encodes several proteins with homology to animal ECM domains, such as laminin G, fibronectin type III, and collagen domains (*40*). Defining the contribution of these proteins to *Capsaspora* aggregation will shed further light into the role of cell-ECM interactions in the emergence of animal multicellularity.

We observed that coDystrophin mutant cells exhibit cortical blebbing. An explanation for this observation that aligns with knowledge of the mammalian DGC is that coDystrophin binds cortical actin and the transmembrane protein coDystroglycan, establishing a coupling across the plasma membrane that increases the structural integrity of the cortex. Loss of coDystrophin would then promote blebbing by reducing this integrity, just as mutation of mammalian dystrophin increases the propensity of the sarcolemma to disassociate from myofibrils in muscle (*41*). However, despite the fact that coDystrophin and coDystroglycan encode the protein domains that mediate their mutual interaction in mammals (WW domain and c-terminal PPxY motif, respectively), the degree of direct interaction of these proteins remains an open question. Further, the putative extracellular ligand or ligands of coDystroglycan are unknown. In muscle, Dystroglycans bind to the laminin G domain present in laminins, which in turn link to extracellular matrix proteins such as collagens, thereby connecting the intracellular cytoskeleton to the ECM mechanically (*1, 42*). The *Capsaspora* genome encodes four predicted secreted proteins encoding laminin G domains and one transmembrane protein encoding laminin G domains (*40, 43*). In fact, this transmembrane protein (CAOG_00286) is among our candidate coYki direct targets (Fig. 1). Determining whether any of these proteins bind coDystroglycan, and whether they contribute to cortical integrity or aggregate morphology, will be important for understanding how an ancestral DGC-like linkage functioned before the evolution of animal extracellular matrices.

Despite evidence that upstream Hippo pathway kinases regulate the subcellular localization of coYki (*24*), the physiological upstream signals that regulate coYki activity remain unknown. Hippo signaling has an established role in animals as a mechanosensor, responding to local forces and substrate stiffness to modulate Yorkie/YAP/TAZ activity (*44*). *Capsaspora* cells are motile but retain a consistent spherical cortical structure that contrasts with the frequent cortical remodeling of amoeboid cells such as neutrophils (*12*). This suggests that mechanisms that stabilize the cortex may be important for *Capsaspora* cell physiology. One possibility is that Hippo signaling contributes to this stability by tuning expression of DGC components, thereby reinforcing membrane–cortex coupling and preventing local cortical deformation or membrane–cortex detachment caused by external forces or actomyosin-derived cortical tension. Testing whether coYki activity responds to mechanical perturbation will be an important direction for future work.

Together, our findings in *Capsaspora* indicate that the DGC emerged before the origin of animal multicellularity and that it was part of an ancient genetic module regulated by Hippo signaling that controlled the stability of the cortical cytoskeleton. Although the full characteristics of the unicellular ancestor of animals remain speculative (*45, 46*), our finding that genetic perturbation of either the *Capsaspora* Hippo pathway or the DGC affects multicellular morphology suggests that this module may have contributed to the regulation of simple multicellular organization in this ancestor. In animal evolution, this function likely underwent modifications due to changes in dystrophin domain architecture, diversification of extracellular ligands, and evolution of tissue-specific expression, eventually giving rise to a specialized role for the DGC in muscle integrity. These studies thus reveal that this protein complex with a key role in human muscle disease has deep, pre-metazoan evolutionary roots in the regulation of cell-surface and cortical mechanical integrity.

## METHODS

### Cell Culture

*Capsaspora* cells were grown in ATCC medium 1034 as described previously (*12*). Staining of *Capsaspora* cells with phalloidin was done as described previously (*47*). Induction of cell aggregation and aggregation in the presence of blebbistatin was done as described previously (*12*). All images of cell aggregates are at 3 days after induction of aggregation.

### Molecular Biology

To generate a vector for stable expression of mScarlet-coDystrophin, we synthesized a gene coding for the coDystrophin amino acid sequence. To minimize the chance of recombination with the endogenous *coDystrophin* gene, we recoded this synthesized gene using synonymous substitutions (*12*), resulting in a transgene with 77 percent identity to the endogenous *coDystrophin* gene at the nucleotide level. This synthetic *coDystrophin* transgene was inserted at the AflII site of pjp103 (*12*) by Gibson assembly, generating a plasmid encoding *mScarlet*-*coDystrophin* and Hygromycin resistance (pJP167). pJP167 sequence is available in DATA.

### Analysis of *Capsaspora* mutant RNAseq data, putative direct coYki targets, and human orthologs

To identify genes that are significantly decreased in the *coYki -/-* mutant, increased in the *coHpo -/-* mutant, and increased in the *coWts -/-* mutant, we examined our previously reported lists of genes differentially expressed in the *coYki* mutant (Supplementary File 1 from (*12*)) and genes differentially expressed in the coHpo and coWts mutants (Supplementary File 2 from (*24*)). To identify putative human orthologs of the nine putative coYki direct targets, we used our previous list of *Capsaspora* genes annotated with putative human orthologs (*12*) as well as annotations from the Uniprot Universal Protein Knowledgebase (*25*).

### *Capsaspora* transfection and genome editing

*Capsaspora* cells were transfected using the TransIT-X2 transfection reagent (Mirus Bio) as previously described (*12*). Stable transgenic cell lines were generated using selection with hygromycin B (Sigma) at 100 μg/mL, geneticin (Thermo Fisher) at 60 μg/mL, or nourseothricin (GoldBio) at 75 μg/mL followed by clonal dilution of drug-resistant cells as described previously (*12*). Homozygous knockout *Capsaspora* cell lines were generated using a gene targeting strategy as described previously (*12*). Briefly, gene targeting constructs consisting of 5’ and 3’ 90 bp segments of homology to the genomic region targeted for deletion and either the Geneticin resistance cassette (GenR) or nourseothricin resistance cassette (NatR) were generated by PCR. Since both of these drug resistance cassettes use the *Capsaspora* actin (CAOG_06018) promoter and terminator, the same primer set can be used to amplify either drug resistance cassette. Plasmids coJP72 and coJP102 (*12*) were used as templates to amplify the GenR and NatR cassette, respectively. WT cells were first transfected with the GenR gene targeting construct, and drug resistant clones were obtained by geneticin selection. Clones with heterozygous mutation of the targeted gene were identified by diagnostic PCR. *Capsaspora* genomic DNA was prepared for diagnostic PCR as described previously (*12*).

These heterozygous mutants were then transfected with the NatR gene-targeting construct, and geneticin + nourseothricin-resistant clones were isolated by drug selection and clonal dilution. Homozygous mutants were then identified by diagnostic PCR (Extended Data Fig. 1). Oligonucleotides used to generate gene targeting constructs are given in DATA, and Oligonucleotides used for diagnostic PCR are given in DATA.

### Microscopy

Confocal imaging of live cells was done using a Zeiss LSM880 microscope. 5 μl of *Capsaspora* cells at 5 x 10^6^ cells/ml in growth medium were added as a droplet at the center of a 4-well glass-bottom chambered coverglass slide (Nunc). Cells were allowed to adhere for 45 minutes, and then 1 ml of growth medium was added to the well, generating a dot of adhered cells in the center of the well. Cells at the edge of the dot were imaged two days after inoculation.

### Measurement of aggregate properties

Images of cell aggregates were manually traced using the ImageJ freehand selection tool. The “measure” function of the ROI manager tool was then used to calculate aggregate area and circularity. To calculate non-circularity, the formula (1-circularity) was used. Three-dimensional aggregate curvature was quantified using Kappa (*48*) as previously described (*12*), with the modification that 4-well borosilicate-bottom chamber slides (Nunc) were used for cell aggregation and imaging. To prepare low-adherence 4-well chamber slides for this protocol, slides were coated with agarose: a 1% molten agarose solution was prepared by microwaving until the agarose was dissolved, 1 mL of molten agarose was added to each well for 5 seconds, the solution was pipetted out of the well and discarded, and the wells were allowed to dry.

### Analysis of dystrophin-like proteins from unicellular organisms and early-branching animals and a *Capsaspora* dystroglycan ortholog

Dystrophin-like proteins were identified by performing BLAST searches using the human dystrophin protein sequence (isoform P11532) against the non-redundant protein sequences database for the following taxonomic groups: Dictyostelia (taxid:33083), Fungi (taxid:4751), Creolimax (taxid:1040653), Choanoflagellata (taxid:28009), Ctenophora (taxid:10197), sponges (taxid:6040), and Nematostella (taxid:45350). Protein IDs for dystrophin-like proteins in each species are: *Capsaspora owczarkaki:* CAOG_003619, *Monosiga brevicollis:* MONBRDRAFT_22129, *Sappingoeca rosetta:* PTSG_05018, *Bolinopsis microptera:* XP_063681244.1, *Mnemiopsis leidyi:* KAL5254423.1, *Sycon ciliatum:* XP_065189684.1 and XP_065185233.1, *Nematostella vectensis:* XP_032238201.2, *Drosophila melanogaster:* Q9VDW6, *Homo sapiens:* P11532, P46939, and Q13474. For both dystrophin-like and dystroglycan-like proteins, protein domains were identified using InterProScan (*49*), NCBI Conserved Domain Search (*50*), or the Uniprot Universal Protein Knowledgebase (*25*). Protein alignments were made using Clustal Omega Multiple Sequence Alignment (*51*). Species images were obtained from PhyloPic (phylopic.org).

### Gene nomenclature

We use unitalicized text to denote a protein (e.g. coDystrophin), italicized text to denote a gene (e.g. *coDystrophin),* “-/-“ to denote a homozygous deletion mutant (e.g. *coDystrophin -/-)*, and “>” to indicate expression of a transgene by a promoter (e.g. *EF1 > coDystrophin*).

### Statistics

Statistical analyses and graphing were done using Prism (GraphPad Software, San Diego, CA). For experiments where multiple cells or aggregates were measured for each independent experiment, a mean was calculated for each independent experiment, and statistical analyses were performed on these biological-replicate means following (*52*). All statistical tests were two-tailed.

## Data Availability

The data that support the findings of this study are available from the corresponding author upon reasonable request.

## Author Contributions

JEP and DP conceived the study. JEP performed the experiments. JEP and DP analyzed the data and wrote the manuscript.

---

**Extended Data Fig. 1:**
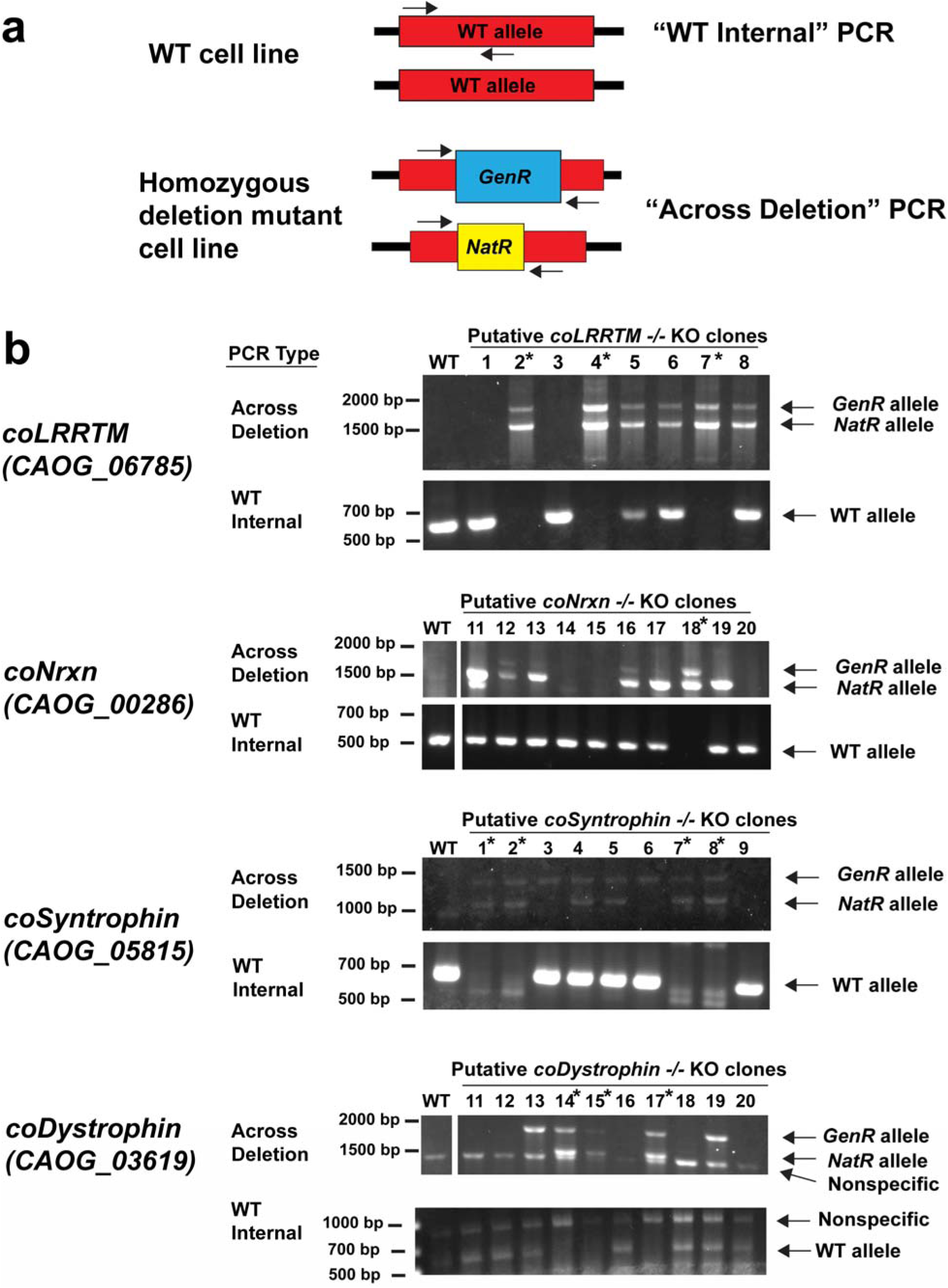
Confirmation of homozygous knockout cell lines by diagnostic PCR. a, Diagrams of diagnostic PCR strategy used to confirm homozygous gene knockout. “WT Internal” PCR will generate a band if a targeted WT genomic sequence is present, but this PCR product will be absent in a homozygous deletion mutant. “Across deletion” PCR will generate bands of unique size if Geneticin resistance cassette (GenR) or Nourseothricin resistance cassette (NatR) are present at the deletion site. Presence of PCR products indicative of both GenR and NatR alleles shows that both WT alleles have been disrupted by insertion of these selectable markers. These primer annealing sites are also present in WT cells, but the predicted WT PCR product is too large to be amplified in the used reaction conditions. **b,** “WT internal” and “Across deletion” PCR products are shown for putative homozygous KO clonal lines of the indicated genotype. Primers are unique for each genotype (see Materials and Methods). Asterisks “*” indicate homozygous mutant clones, as confirmed by absence of the WT allele and presence of both GenR and NatR deletion alleles. Some clones show PCR products for all three alleles; these are likely mixed genetic populations.

**Video S1: Visualization of F-actin in adherent *Capsaspora* cells stably expressing Lifeact-mScarlet.** Video is 4 minutes and 19 seconds long. This video serves as a control for Video S2.

**Video S2: Visualization of F-actin in adherent *coDystrophin -/-* cells stably expressing Lifeact-mScarlet.** Video is 4 minutes and 19 seconds long.

**Video S3: Overexpression of *coDystrophin* suppresses blebbing of *coYki* mutant cells.** *coYki -/-* cells were transiently transfected with Lifeact-mNeon to visualize F-actin (green signal) and either mScarlet (left, red signal) or an mScarlet-coDystrophin fusion (right, red signal). Scale bar is 5 microns. Time is shown in minutes:seconds.

**Video S4: Expression of human Dystrophin suppresses blebbing in *coDystrophin* mutant cells.** *coDystrophin* -/- cells were transiently transfected with Lifeact-mNeon to visualize F-actin (green signal) and either mScarlet (left, red signal), mScarlet-coDystrophin (center), or mScarlet-hsDystrophin (right). Time is shown in minutes:seconds.

## REFERENCES

1. J. M. Ervasti, K. P. Campbell, A Role for the Dystrophin-Glycoprotein Complex as a Transmembrane Linker between Laminin and Actin. Journal of Cell Biology 122, 809–823 (1993).

2. A. Suzuki, M. Yoshida, H. Yamamoto, E. Ozawa, Glycoprotein-Binding Site of Dystrophin Is Confined to the Cysteine-Rich Domain and the 1st-Half of the Carboxy-Terminal Domain. Febs Letters 308, 154–160 (1992).

3. O. Ibraghimovbeskrovnaya et al., Primary Structure of Dystrophin-Associated Glycoproteins Linking Dystrophin to the Extracellular-Matrix. Nature 355, 696–702 (1992).

4. F. Muntoni, S. Torelli, A. Ferlini, Dystrophin and mutations: one gene, several proteins, multiple phenotypes. Lancet Neurol 2, 731–740 (2003).

5. K. Tetorou, A. Aghaeipour, S. Singh, J. E. Morgan, F. Muntoni, The role of dystrophin isoforms and interactors in the brain. Brain 148, 1081–1098 (2025).

6. Q. Q. Gao, E. M. McNally, The Dystrophin Complex: Structure, Function, and Implications for Therapy. Compr Physiol 5, 1223–1239 (2015).

7. S. Le et al., Dystrophin As a Molecular Shock Absorber. ACS Nano 12, 12140–12148 (2018).

8. D. S. Duan, N. Goemans, S. Takeda, E. Mercuri, A. Aartsma-Rus, Duchenne muscular dystrophy. Nat Rev Dis Primers 7, (2021).

9. J. C. Adams, A. Brancaccio, The evolution of the dystroglycan complex, a major mediator of muscle integrity. Biol Open 4, 1163–1179 (2015).

10. D. T. Schultz et al., Ancient gene linkages support ctenophores as sister to other animals. Nature 618, 110–117 (2023).

11. V. Mirouse, Evolution and developmental functions of the dystrophin-associated protein complex: beyond the idea of a muscle-specific cell adhesion complex. Front Cell Dev Biol 11, 1182524 (2023).

12. J. E. Phillips, M. Santos, M. Konchwala, C. Xing, D. Pan, Genome editing in the unicellular holozoan Capsaspora owczarzaki suggests a premetazoan role for the Hippo pathway in multicellular morphogenesis. Elife 11, (2022).

13. A. Sebe-Pedros, A. J. Roger, F. B. Lang, N. King, I. Ruiz-Trillo, Ancient origin of the integrin-mediated adhesion and signaling machinery. Proc Natl Acad Sci U S A 107, 10142–10147 (2010).

14. H. Suga et al., Genomic survey of premetazoans shows deep conservation of cytoplasmic tyrosine kinases and multiple radiations of receptor tyrosine kinases. Sci Signal 5, ra35 (2012).

15. A. Sebe-Pedros, A. de Mendoza, B. F. Lang, B. M. Degnan, I. Ruiz-Trillo, Unexpected repertoire of metazoan transcription factors in the unicellular holozoan Capsaspora owczarzaki. Mol Biol Evol 28, 1241–1254 (2011).

16. S. A. Nichols, B. W. Roberts, D. J. Richter, S. R. Fairclough, N. King, Origin of metazoan cadherin diversity and the antiquity of the classical cadherin/beta-catenin complex. Proc Natl Acad Sci U S A 109, 13046–13051 (2012).

17. N. Ros-Rocher et al., Chemical factors induce aggregative multicellularity in a close unicellular relative of animals. Proc Natl Acad Sci U S A 120, e2216668120 (2023).

18. A. Sebe-Pedros, Y. Zheng, I. Ruiz-Trillo, D. Pan, Premetazoan origin of the hippo signaling pathway. Cell Rep 1, 13–20 (2012).

19. J. E. Phillips, Y. Zheng, D. Pan, Assembling a Hippo: the evolutionary emergence of an animal developmental signaling pathway. Trends Biochem Sci, (2024).

20. D. Pan, The unfolding of the Hippo signaling pathway. Dev Biol 487, 1–9 (2022).

21. G. Halder, R. L. Johnson, Hippo signaling: growth control and beyond. Development 138, 9–22 (2011).

22. Z. P. Meng, T. Moroishi, K. L. Guan, Mechanisms of Hippo pathway regulation. Gene Dev 30, 1–17 (2016).

23. T. Panciera, L. Azzolin, M. Cordenonsi, S. Piccolo, Mechanobiology of YAP and TAZ in physiology and disease. Nat Rev Mol Cell Biol 18, 758–770 (2017).

24. J. E. Phillips, D. Pan, The Hippo kinase cascade regulates a contractile cell behavior and cell density in a close unicellular relative of animals. Elife 12, (2024).

25. C. UniProt, UniProt: the Universal Protein Knowledgebase in 2025. Nucleic Acids Res 53, D609–D617 (2025).

26. I. Diamant, D. J. B. Clarke, J. E. Evangelista, N. Lingam, A. Ma’ayan, Harmonizome 3.0: integrated knowledge about genes and proteins from diverse multi-omics resources. Nucleic Acids Res 53, D1016–D1028 (2025).

27. S. C. Watkins, E. P. Hoffman, H. S. Slayter, L. M. Kunkel, Immunoelectron microscopic localization of dystrophin in myofibres. Nature 333, 863–866 (1988).

28. R. G. Roberts, Dystrophins and dystrobrevins. Genome Biol 2, REVIEWS3006 (2001).

29. D. Jung, B. Yang, J. Meyer, J. S. Chamberlain, K. P. Campbell, Identification and characterization of the dystrophin anchoring site on beta-dystroglycan. J Biol Chem 270, 27305–27310 (1995).

30. M. James et al., Adhesion-dependent tyrosine phosphorylation of (beta)-dystroglycan regulates its interaction with utrophin. J Cell Sci 113 ( Pt 10), 1717–1726 (2000).

31. A. Akhavan, S. N. Crivelli, M. Singh, V. R. Lingappa, J. L. Muschler, SEA domain proteolysis determines the functional composition of dystroglycan. FASEB J 22, 612–621 (2008).

32. K. A. Deyst, M. A. Bowe, J. D. Leszyk, J. R. Fallon, The alpha-dystroglycan-beta-dystroglycan complex. Membrane organization and relationship to an agrin receptor. J Biol Chem 270, 25956–25959 (1995).

33. E. Kimura et al., Cell-lineage regulated myogenesis for dystrophin replacement: a novel therapeutic approach for treatment of muscular dystrophy. Human Molecular Genetics 17, 2507–2517 (2008).

34. I. Lian et al., The role of YAP transcription coactivator in regulating stem cell self-renewal and differentiation. Genes Dev 24, 1106–1118 (2010).

35. Y. Morikawa et al., Actin cytoskeletal remodeling with protrusion formation is essential for heart regeneration in Hippo-deficient mice. Sci Signal 8, ra41 (2015).

36. Y. Morikawa, T. Heallen, J. Leach, Y. Xiao, J. F. Martin, Dystrophin-glycoprotein complex sequesters Yap to inhibit cardiomyocyte proliferation. Nature 547, 227–231 (2017).

37. C. Nielsen, Early animal evolution: a morphologist’s view. R Soc Open Sci 6, 190638 (2019).

38. S. Rentschler et al., The WW domain of dystrophin requires EF-hands region to interact with beta-dystroglycan. Biol Chem 380, 431–442 (1999).

39. T. C. Levin, A. J. Greaney, L. Wetzel, N. King, The Rosetteless gene controls development in the choanoflagellate S. rosetta. Elife 3, (2014).

40. F. Williams, H. A. Tew, C. E. Paul, J. C. Adams, The predicted secretomes of Monosiga brevicollis and Capsaspora owczarzaki, close unicellular relatives of metazoans, reveal new insights into the evolution of the metazoan extracellular matrix. Matrix Biol 37, 60–68 (2014).

41. K. P. Garcia-Pelagio, R. J. Bloch, A. Ortega, H. Gonzalez-Serratos, Biomechanics of the sarcolemma and costameres in single skeletal muscle fibers from normal and dystrophin-null mice. J Muscle Res Cell Motil 31, 323–336 (2011).

42. J. F. Talts, Z. Andac, W. Göhring, A. Brancaccio, R. Timpl, Binding of the G domains of laminin α1 and α2 chains and perlecan to heparin, sulfatides, α-dystroglycan and several extracellular matrix proteins. Embo Journal 18, 863–870 (1999).

43. H. Suga et al., The Capsaspora genome reveals a complex unicellular prehistory of animals. Nat Commun 4, 2325 (2013).

44. B. K. Tripathi, K. D. Irvine, Mechanobiology of the Hippo-YAP Signaling Network. Cold Spring Harb Perspect Biol, (2026).

45. N. Ros-Rocher, A. Perez-Posada, M. M. Leger, I. Ruiz-Trillo, The origin of animals: an ancestral reconstruction of the unicellular-to-multicellular transition. Open Biol 11, 200359 (2021).

46. T. Brunet, N. King, The Origin of Animal Multicellularity and Cell Differentiation. Dev Cell 43, 124–140 (2017).

47. H. Parra-Acero et al., Integrin-Mediated Adhesion in the Unicellular Holozoan Capsaspora owczarzaki. Curr Biol 30, 4270–4275 e4274 (2020).

48. H. Mary, G. J. Brouhard, Kappa (κ): Analysis of Curvature in Biological Image Data using B-splines. bioRxiv, 852772 (2019).

49. P. Jones et al., InterProScan 5: genome-scale protein function classification. Bioinformatics 30, 1236–1240 (2014).

50. J. Wang et al., The conserved domain database in 2023. Nucleic Acids Res 51, D384–D388 (2023).

51. F. Sievers et al., Fast, scalable generation of high-quality protein multiple sequence alignments using Clustal Omega. Mol Syst Biol 7, 539 (2011).

52. S. J. Lord, K. B. Velle, R. D. Mullins, L. K. Fritz-Laylin, SuperPlots: Communicating reproducibility and variability in cell biology. J Cell Biol 219, (2020).

53. H. Parra-Acero et al., Transfection of Capsaspora owczarzaki, a close unicellular relative of animals. Development 145, (2018).

